# Low-cost, handheld near-infrared spectroscopy for root dry matter content prediction in cassava

**DOI:** 10.1101/2021.04.30.441802

**Authors:** Jenna Hershberger, Edwige Gaby Nkouaya Mbanjo, Prasad Peteti, Andrew Smith Ikpan, Kayode Ogunpaimo, Kehinde Nafiu, Ismail Y. Rabbi, Michael A. Gore

## Abstract

Over 800 million people across the tropics rely on cassava as a major source of calories. While the root dry matter content (RDMC) of this starchy root crop is important for both producers and consumers, characterization of RDMC by traditional methods is time-consuming and laborious for breeding programs. Alternate phenotyping methods have been proposed but lack the accuracy, cost, or speed ultimately needed for cassava breeding programs. For this reason, we investigated the use of a low-cost, handheld NIR spectrometer for field-based RDMC prediction in cassava. Oven-dried measurements of RDMC were paired with 21,044 scans of roots of 376 diverse clones from 10 field trials in Nigeria and grouped into training and test sets based on cross-validation schemes relevant to plant breeding programs. Mean partial least squares regression model performance ranged from R^2^_p_ = 0.62 - 0.89 for within-trial predictions, which is within the range achieved with laboratory-grade spectrometers in previous studies. Relative to other factors, model performance was highly impacted by the inclusion of samples from the same environment in both the training and test sets. Random forest variable importance analysis of root spectra revealed increased importance in a region previously identified as predictive of water content in plants (~950 - 990 nm). With appropriate model calibration, the tested spectrometer will allow for field-based collection of spectral data with a smartphone for accurate RDMC prediction and potentially other quality traits, a step that could be easily integrated into existing harvesting workflows of cassava breeding programs.

**CORE IDEAS:**

- A low-cost, handheld near-infrared spectrometer was tested for phenotyping of cassava roots
- Plant breeding-relevant cross-validation schemes were used for predictions
- High prediction accuracies were achieved for cassava root dry matter content
- A spectral region predictive of plant water content was identified as important

## INTRODUCTION

Cassava is a starch-rich industrial root crop and staple food for hundreds of millions of people throughout the tropics (Howeler et al., 2013). Although it is especially valued for its role as a subsistence crop, as it is able to withstand environmental conditions that other crops cannot and can be relied upon in times of drought and food scarcity (El-Sharkawy, 1993, 2004), its role in commercial farming and industrial processing is increasing globally (Parmar et al., 2017). Though more than half of global cassava production occurs in Africa, yields on the continent remain far below the global average (FAO, 2021). While a portion of this yield gap is likely due to management factors such as differences in fertilizer applications, low rates of improved germplasm adoption also contribute to this gap (Fermont et al., 2009; Njine, 2010; Oparinde et al., 2016; Owusu & Donkor, 2012).

As is true for other crops, this lack of improved variety adoption is largely attributed to the need for breeding objectives to place more weight on the selection for end user preferred traits (Acheampong et al., 2018; Asrat et al., 2010). Early breeding objectives focused heavily on the development of varieties resistant to diseases that threatened to wipe out cassava production across the African continent (Hahn et al., 1980; IITA, 1990; Jennings, 1957), but following success in those efforts, emphasis has since shifted to breeding for yield and culturally-relevant quality traits. Root dry matter content (RDMC), a major component of both dry yield and food quality that is of high importance along the entire value chain (Okechukwu & Dixon, 2008; Teeken et al., 2018, 2020), is now directly included in the selection indices used by major cassava breeding programs including those implementing genomic selection (Ceballos et al., 2012; Kawano, 2003; Kawuki et al., 2011; Wolfe et al., 2016).

Traditional methods of cassava RDMC characterization are time-consuming and laborious, and therefore unsuitable for phenotyping the large number of plots at the early stages of selection. Oven drying, the gold standard phenotyping method for RDMC, is not only tedious but also requires a cheap and stable source of heat energy. However, this is not available in the majority of off-station testing locations, thereby necessitating transportation of roots to centralized laboratories (Safo-Kantanka & Owusu-Nipah, 1992; Teles et al., 1993; Teye et al., 2011). To work around these obstacles, researchers introduced a simple linear regression equation to relate specific gravity to RDMC (Kawano et al., 1987), a method that can be performed directly in the field with just a scale and basin of water. However, due to the relatively thick peels of cassava roots and an inability to wash them thoroughly in the field, this method is often inaccurate (Ikeogu et al., 2017; Pérez et al., 2011), limiting its potential benefits.

More recently, near-infrared spectroscopy (NIRS) was introduced to cassava breeding as an alternative method for RDMC phenotyping (Sánchez et al., 2014). With the correct model calibration, NIRS offers highly accurate estimates of RDMC at a fraction of the time of oven drying, potentially increasing the throughput without proportional rises in cost. Sánchez et al. found the Foss 6500, a lab-grade, benchtop visible and near-infrared (Vis-NIR) spectrometer, to be highly predictive (R^2^_p_ = 0.79 - 0.95) of RDMC in cassava over several growing seasons.

While the use of a benchtop spectrometer saves effort compared to oven drying, it still requires root transport and extensive sample preparation, ultimately resulting in an equivalent amount of labor to the oven method of RDMC determination. This study was followed by an evaluation of the ASD QualitySpec Trek (Malvern Panalytical), a mobile Vis-NIR spectrometer, that found it to be sufficiently accurate for RDMC prediction in breeding programs (Ikeogu et al., 2017). Though this instrument can be taken to the field and does not require samples to be shredded or blended before scanning, its high cost limits accessibility for breeding programs with more rigid budget constraints.

To truly meet the needs of cassava breeding programs, there is a need for a spectrometer that is accurate, field-based, and low-cost. Recent developments in NIRS technology have resulted in smaller, light weight devices that require minimal sample preparation, effectively allowing the user to bring the laboratory to the sample (Teixeira Dos Santos et al., 2013). Many of these mobile spectrometers (e.g., ConsumerPhysics: SCiO; Stratio: LinkSquare; Tellspec Inc.: Tellspec; Felix Instruments Inc.: F-750 Produce Quality Meter) are also considerably less expensive than their predecessors, which could facilitate their adoption by breeding programs with limited funding. The application of this technology in a cassava breeding context has the potential to boost the throughput of RDMC phenotyping without increasing the cost or time investment per sample, but to our knowledge, the use of lower cost, mobile NIR spectrometers have not been reported for quantification of RDMC in cassava.

In this study, we developed and evaluated a new phenotyping procedure to predict cassava RDMC in an active breeding program. The main objectives of the study were to (i) assess a low-cost, handheld NIR spectrometer for the prediction of cassava RDMC, and (ii) evaluate the utility of collected NIRS data in the context of a cassava breeding program, developing best practices for routine use.

## MATERIALS AND METHODS

### Plant materials and experimental design

In order to test spectrometer and model performance within and across populations and environments, 10 field experiments were evaluated (Table 1). These experiments were representative of field trials commonly used to test the genetic potential of cassava clones for a range of phenotypes, containing clones of varying levels of improvement, relatedness, and RDMC. A complete list of clones and corresponding phenotypic data are available on Cassavabase (www.cassavabase.org).

**Table 1.** Metadata for 10 IITA cassava field trials

| Abbreviated Trial Name | Cassavabase Trial Name | Trial Type<sup>a</sup> | Planting Date | Harvest Date | Experimental design<sup>b</sup> | # Replicates | Total # unique clones planted | # Check clones planted | # Plants per plot | Plot dimensions (m) |
| --- | --- | --- | --- | --- | --- | --- | --- | --- | --- | --- |
| A-17IB | 17.CASS.PY T.49.setA.IB | PYT | 2017-Apr.-21 | 2018-May-09 | RCBD | 2 | 49 | 4 | 10 | 2 x 4 |
| B-17IB | 17.GS.C3.PY T.80.IB | PYT | 2017-May-11 | 2018-May-17 | Alpha-lattice | 2 | 80 | 6 | 10 | 2 x 4 |
| C-18IB | 18.CASS.PY T.52.IB | PYT | 2018-July-05 | 2019-July-05 | RCBD | 2 | 52 | 4 | 36 | 6 x 4.8 |
| D-18IB | 18.GS.C2.set A.UYT.36.IB | UYT | 2018-July-19 | 2019-July-05 | Alpha-lattice | 3 | 36 | 5 | 42 | 6 x 5.6 |
| E-18IB | 18.GS.C2.set B.UYT.36.IB | UYT | 2018-Jan.-21 | 2019-July-19 | Alpha-lattice | 3 | 36 | 5 | 42 | 6 x 5.6 |
| F-19IB | 19.CMSSurveyVarieties.A YT.33.IB | AYT | 2019-Apr.-29 | 2020-Apr.-20 | RCBD | 2 | 33 | 5 | 20 | 4 x 4 |
| G-19IB | 19.GS.C2.UYT.36.setA.IB | UYT | 2019-May-16 | 2020-Apr.-20 | RCBD | 2 | 36 | 5 | 20 | 4 x 4 |
| H-19IB | 19.GS.C2.UYT.36.setB.IB | UYT | 2019-May-16 | 2020-Apr.-28 | RCBD | 2 | 36 | 5 | 20 | 4 x 4 |
| I-19IK | 19.CASS.PY T.52.IK | PYT | 2019-June-25 | 2020-July-20 | RCBD | 2 | 52 | 4 | 20 | 4 x 4 |
| J-19IK | 19.GS.C4B.PYT.500.IK | PYT | 2019-Aug.-04 | 2020-Oct.-27 | Alpha-lattice | 2 | 452 | 5 | 9 | 3 x 2.4 |

<sup>a</sup>AYT = Advanced yield trial; PYT = Preliminary yield trial; UYT = Uniform yield trial
<sup>b</sup>RCBD = randomized complete block design

Eight of the field trials were planted at the International Institute for Tropical Agriculture (IITA) in Ibadan, Nigeria (A-17IB, B-17IB, C-18IB, D-18IB, E-18IB, F-19IB, G-19IB, H-19IB), while two were planted in Ikenne, Nigeria (I-19IK and J-19IK). All trials were planted with 1 m inter-row spacing and 0.8 m alleys at the end of each plot. Phenotypic data were collected on two replicates of each experimental clone in each trial.

A-17IB, C-18IB, and I-19IK included clones selected from the genetic gain germplasm collection, a population of historically important cassava clones from IITA, as previously described by Okechukwu and Dixon (2008), Ly et al. (2013), and Wolfe et al. (2016). F-19IB included clones sampled from a survey of popular varieties in Nigeria. Clones in all other trials were derived from various cycles of genomic selection originating from the genetic gain population: trials D-18IB, E-18IB, G-19IB and H-19IB consisted of clones from the second cycle, B-17IB clones were selected from the third cycle, and J-19IK included clones from the fourth cycle.

### Dry matter determination and spectral data acquisition

The SCiO molecular sensor (Consumer Physics Inc., Tel-Aviv, Israel) is a handheld, portable NIR spectrometer that connects to the SCiO Lab smartphone application via Bluetooth for spectral data acquisition. Using an active light source, it measures diffuse reflectance from 740-1070 nm, providing values in 1 nm intervals. Each of these measurement scans takes approximately two seconds and is immediately transferred to the SCiO server for data storage. Prior to each use, calibration was performed using the built-in reference standard in the SCiO case. A manufacturer-provided plastic “light shield” was attached to the spectrometer for all scans to block ambient light and to maintain a consistent 9 mm distance from the sensor to the sample surface during the scanning process.

Six commercial-sized roots were collected on a plot basis for all trials. In A-17IB, B-17IB, and C-18IB, roots were sliced in half and scanned with the SCiO three times on each cut surface for a total of six scans per root and 36 scans per plot (Figure 1, Supplementary Figure 1). After scanning, individual roots were peeled, shredded and a sample of 100 g was oven-dried at 80°C until reaching a constant weight (48 hr). Percent RDMC was calculated as:

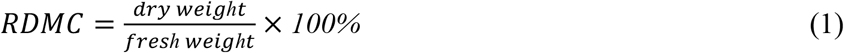

**Figure 1.**
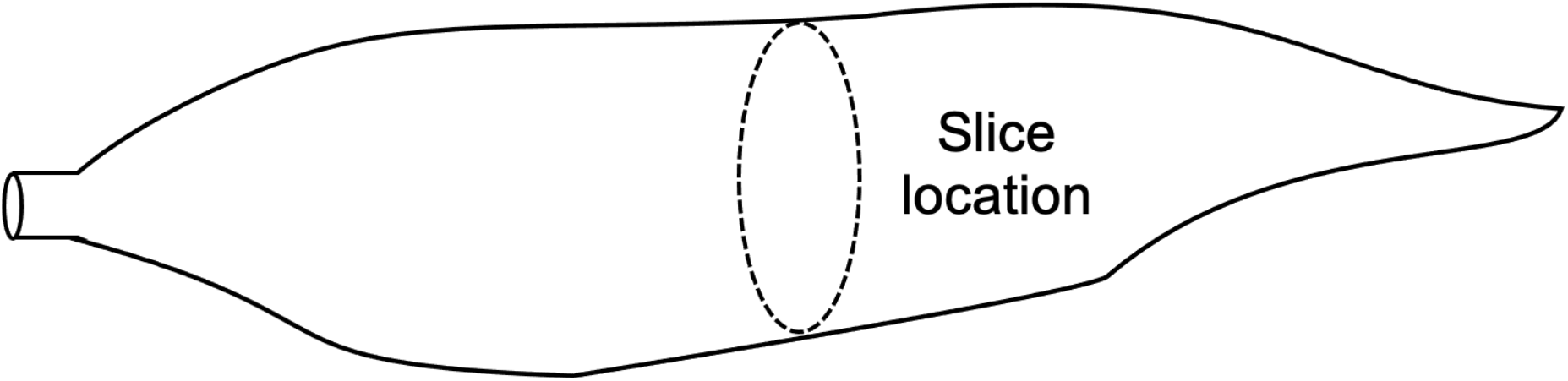
Roots from trials A-17IB, B-17IB, and C-18IB were sliced crosswise in the central region as shown and scans were taken on each side of the cut surface.

In the seven remaining trials, scans were taken after grating the root to obtain a homogeneous plot-level sample. Three to ten subsamples of homogenized fresh root tissue were placed in a quartz glass container and scanned. Percent RDMC was obtained using the same oven-drying method as the other three trials. Individual field plots with incomplete spectral or RDMC data were removed from the analysis, at times resulting in a single replicate of a given clone in the final dataset.

### Outlier removal and sample aggregation

After data collection, all scans were filtered according to Mahalanobis distance. Samples with Mahalanobis distances greater than a cutoff set by a χ^2^-distribution with 331 degrees of freedom (*α* = 0.05) were removed from the analysis (Johnson & Wichern, 2007) using the *waves* R package version 0.1.1 (Hershberger et al., 2021) in R version 3.5.2 (R Core Team, 2018). In total, eight scans were removed through this procedure. These outlier scans are listed in Supplemental Table S1. After outlier removal, sample aggregation by means, as typical for NIRS studies (Ikeogu et al., 2017; A. Kaur et al., 2020; Lebot et al., 2009; Sánchez et al., 2014), was performed to enable RDMC prediction on a plot-level basis (i.e., the experimental unit) for all trials.

### Spectral preprocessing

Twelve combinations of common preprocessing methods including standard normal variate (Barnes et al., 1989), first and second derivatives, and Savitzky-Golay polynomial smoothing (Savitzky & Golay, 1964) were applied using the R package *waves* version 0.1.1 (Hershberger et al., 2021). No clear differences in model performance were found between raw and preprocessed data using any of the preprocessing methods for within-trial predictions (Supplemental Figure S2, Supplemental Table S2), thus raw data were used for all subsequent analyses.

### Cross-validation

A cross-validation scheme was developed for within-trial predictions. In this scheme, samples were separated into training and test sets (70%: 30%) using stratified random sampling to ensure that the full range of RDMC values was represented in each set. This scheme did not take experimental or check plots into account, so in some cases different plots from the same experimental or check clones were included in both the training and test sets.

Four additional cross-validation (CV) schemes representing relevant plant breeding scenarios were applied across the 10 trials with each trial considered an independent environment due to differences in planting and harvest dates even within the same year. These CV schemes are described in detail in Jarquín et al. (2017), but in brief, all pairwise combinations of tested and untested genotypes in tested and untested environments were evaluated in four CV schemes (Figure 2). For CV2 (tested genotypes in tested environments; Figure 2A), a random subset of 30% of the clones from a given trial were used as the test set. Data from all remaining clones from that trial and all other trials were combined into one training set. This process was repeated with 50 iterations for each test trial, each with a different random sample of clones in the test set. CV1 (untested genotypes in tested environments; Figure 2B) was implemented using the same subsetting procedure as CV2. In this scheme, however, clones present in the test set were removed from the training set altogether. CV0 (tested genotypes in untested environments; Figure 2C) involved the inclusion of an entire trial as the test set, with all other trials making up the training set whether or not they contained clones represented in the test set trial. The implementation of CV00 (untested genotypes in untested environments; Figure 2D) followed the same procedure as CV0, but as with CV1, all clones present in the test set were removed from the training set prior to model training. In all cases, all plots of each clone within a given trial were grouped, occurring either both in the training or both in the test set. Counts of overlapping clones between trials are listed in Supplemental Table S3. All CV schemes were implemented in the *waves* R package version 0.1.1 (Hershberger et al., 2021).

**Figure 2.**
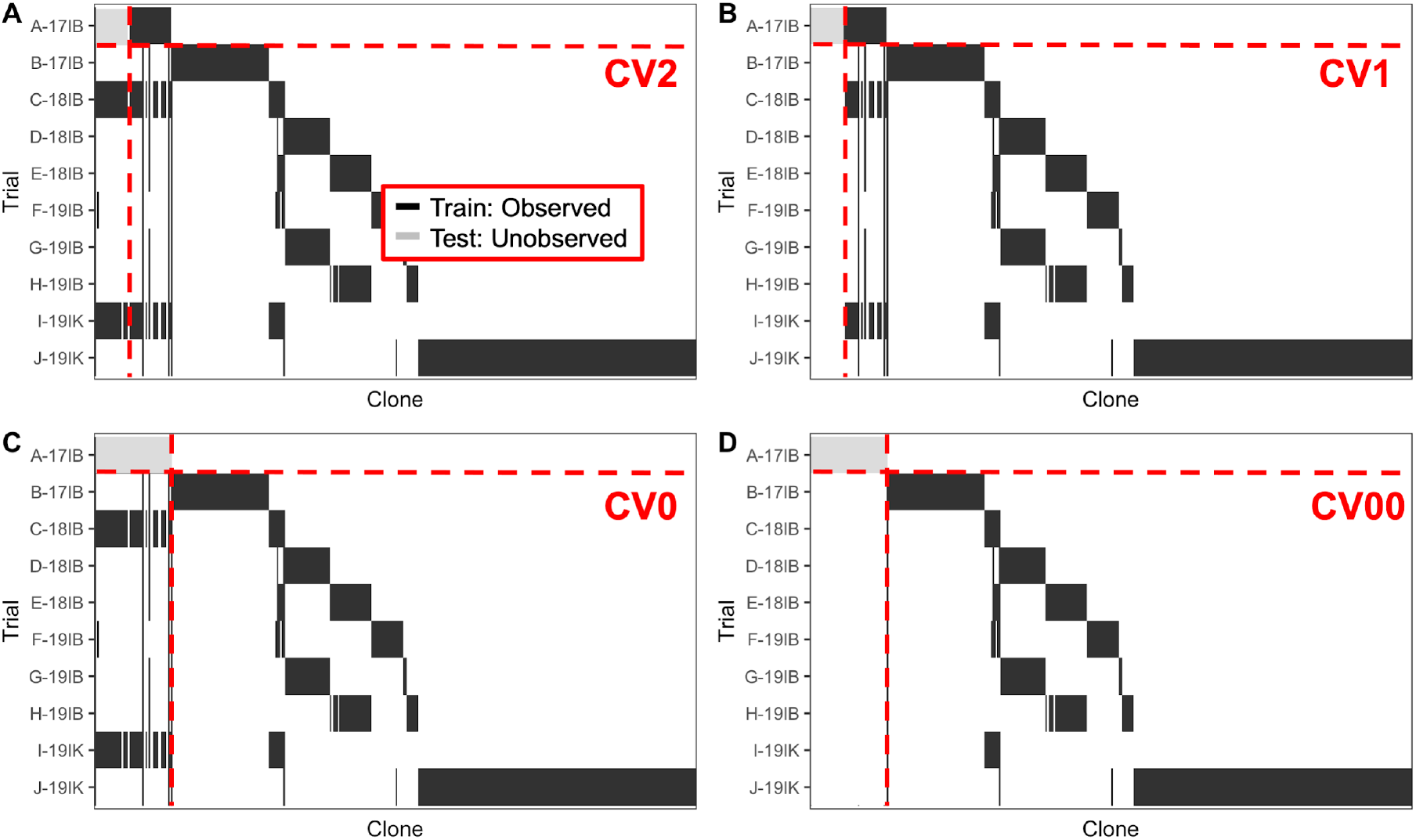
Graphic representation of four of the five cross-validation (CV) schemes used in this study. Unique cassava clones are shown along the x-axis and individual trials are along the y-axis. Clones appearing in test sets are represented in gray, while clones in the training sets are black. In this example, various proportions of clones from trial A-17IB are included in the test set according to the cross-validation scheme. (a) CV2 represents a case in which observations of the test set clones are present in the training set along with observations of clones from all environments. (b) CV1 also includes observations from all environments, but no observations from the test set clones are included in the training set. (c) CV0 includes observations from all clones but includes all clones from trial A-17IB in the test set so that environment is not represented in the training set. (d) CV00 omits all observations from the training set that overlap with the test set in clone or environment. These CV schemes are modeled after Jarquín et al. (2017).

### Prediction

Prediction models were developed using three different algorithms in the *waves* R package version 0.1.1 (Hershberger et al., 2021): partial least squares regression (PLSR) (H. Wold, 1982; S. Wold et al., 1984), random forest (RF) (Breiman, 2001), and support vector machine (SVM) (Drucker et al., 1997). PLSR both displayed the best performance for within-trial predictions and required the least time-intensive training (Supplemental Table S2), so it was used for all subsequent prediction analyses in this study.

PLSR is a popular method used in NIRS analysis (reviewed in Roggo et al., 2007). This algorithm uses principal components to reduce the dimensionality of the spectral dataset, allowing for efficient model training. For each CV scenario, the number of components used in the final model was tuned using 5-fold CV within the training set. The best number of components was chosen based on the lowest root mean squared error of CV (RMSE_cv_) and predictions were made on the test set using this final tuned model. This process of sampling and tuning was repeated 50 times for each combination of preprocessing technique and algorithm for a total of 1,950 runs per trial. For CV0 and CV00 scenarios, only a single iteration of hyperparameter tuning and prediction was performed, as no sampling was performed in these cases.

This pipeline was also used to evaluate the optimal number of scanned samples (homogenized subsamples or sliced root subsets) per plot. For trials in which root slices were scanned, A-17IB, B-17IB, and C-18IB, a sample was defined as a single root. For all remaining trials, each 100 g homogenized subsample within a plot was treated as an individual sample. Within each trial, *n* versions of the RDMC and spectral dataset were created, each containing a different number of samples per plot with *n* representing the maximum number of samples available for that trial. For the versions with one to *n*-1 samples per plot, where a different individual or combination of individual samples may be selected from each plot, 1000 iterations of random statistical subsampling were performed to get a representative sample of the possible combinations of homogenized subsamples or sliced root subsets. Each version of the dataset was then run through the *waves* within-trial tuning and prediction pipeline using PLSR for model performance comparison.

### Variable importance

To facilitate biological interpretation, plot mean spectra without preprocessing were used to calculate the random forest variable importance measure for each wavelength within each trial (Breiman, 2001). In short, the prediction accuracy was calculated using the entire spectral data training set and then repeated with each wavelength individually withheld. The difference between model performance in each permutation was then averaged over all decision trees and normalized by the standard error (Liaw & Wiener, 2002). This procedure was performed 10 times, each with a different beginning training set of 70% of the plots as selected using stratified random sampling based on RDMC. All plots for this and other analyses were prepared with *ggplot2* version 3.3.3 (Wickham, 2016).

## RESULTS AND DISCUSSION

Mobile NIRS has the potential to provide rapid, in-field phenotyping of cassava roots for dry matter content, but validation is required to ensure that realistic expectations can be set regarding the accuracy of prediction with a given spectrometer and statistical model. We explored the use of an inexpensive, handheld NIR spectrometer for the prediction of cassava RDMC in the context of an active breeding program. The results presented here demonstrate the feasibility of RDMC prediction with low-cost, mobile spectrometers such as the SCiO used in our study and inform its use in routine breeding decisions. To the best of our knowledge, this work represents the first use of a consumer-grade spectrometer for RDMC prediction in cassava.

### Variation for dry matter content of cassava roots

We measured the RDMC of 376 cassava clones evaluated in one or more of 10 field trials in Nigeria. Summary statistics of plot averages of raw RDMC values as measured with the oven method for the 10 trials in this study are shown in Table 2, with plot mean RDMC distributions shown in Figure 3. Tukey’s honest significant difference test identified significant differences (*α* = 0.05) between the mean values of almost all trials. I-19-IK contained the largest range of RDMC, spanning from 9.75% to 43.30%, resulting in a standard deviation of 7.86. The two other trials with this same set of germplasm, A-17IB and C-18IB, had the second (6.17) and third (5.69) highest standard deviations. Maximum RDMC ranged from 36.0% in H-19IB to 47.1% in J-19IK, with maximums from the other trials falling in between. Minimum RDMC ranged from 9.75% (I-19IK) to 30.6% (E-18IB). The range of RDMC across the 10 trials (9.75 - 47.10%) is representative of the typical RDMC range in cassava germplasm, with adequate variation for improvement through breeding (Sánchez et al., 2009).

**Table 2.** Root dry matter content (RDMC) summary statistics for 10 IITA cassava field trials. The number of clones and plots includes only those with complete spectral and phenotypic data.

| Abbreviated Trial Name | # Phenotyped experimental/check clones | # Phenotyped Plots | Mean RDMC | Maximum RDMC | Minimum RDMC | DMC Standard Deviation |
| --- | --- | --- | --- | --- | --- | --- |
| A-17IB | 44/4 | 92 | 24.2 | 39.8 | 10.8 | 6.2 |
| B-17IB | 59/6 | 156 | 32.5 | 42.0 | 21.8 | 4.1 |
| C-18IB | 47/4 | 97 | 32.1 | 43.7 | 19.3 | 5.7 |
| D-18IB | 30/5 | 103 | 40.7 | 45.9 | 29.7 | 3.1 |
| E-18IB | 31/5 | 100 | 40.1 | 45.9 | 30.6 | 2.8 |
| F-19IB | 25/5 | 52 | 28.5 | 39.1 | 17.1 | 4.6 |
| G-19IB | 30/5 | 68 | 33.6 | 43.2 | 26.7 | 3.6 |
| H-19IB | 31/5 | 65 | 30.9 | 36.0 | 21.7 | 2.7 |
| I-19IK | 46/4 | 98 | 27.6 | 43.3 | 9.8 | 7.9 |
| J-19IK | 175/5 | 331 | 37.9 | 47.1 | 27.0 | 4.2 |

**Figure 3.**
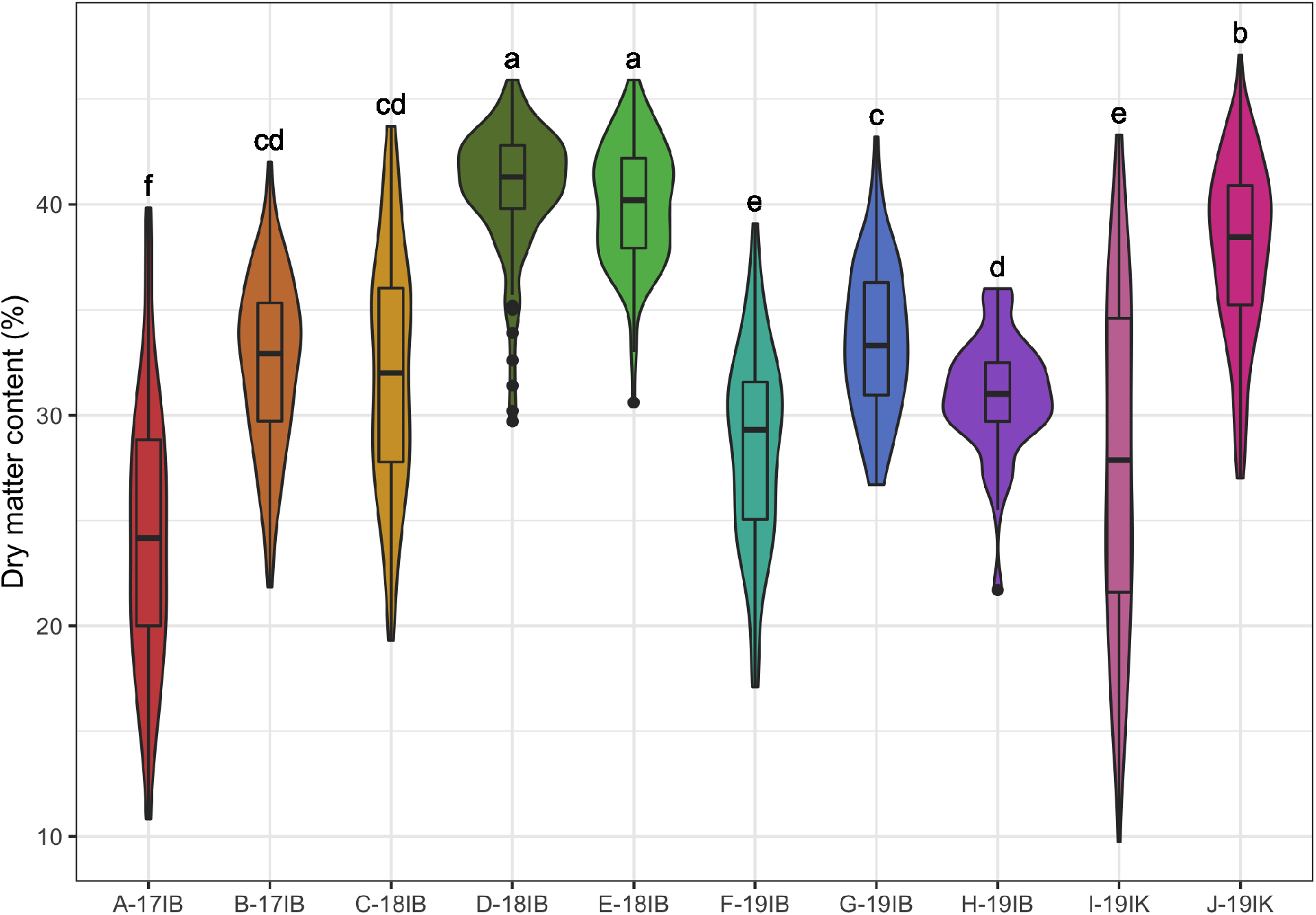
Plot mean root dry matter content distributions are shown for each trial. Letters above the violin and boxplots represent significantly different groups as identified by Tukey’s honest significant different test at *α* = 0.05.

### Within-trial prediction re

To assess within-trial PLSR prediction accuracy of RDMC, we used a stratified random sample of 70% of the plots in a given trial as the training set for hyperparameter tuning and testing model performance on the remaining 30% of plots. Mean prediction performance over 50 iterations of subsampling varied across years, locations, populations, and sample preparation methods (Table 3 and Figure 4), with the highest squared Pearson’s correlation of predicted and observed values from the test set (R^2^_p_) from I-19IK at 0.89 and the lowest root mean squared error of prediction (RMSE_p_) from H-19IB at 1.31. The poorest performing trials in terms of R^2^_p_ were C-18IB and D-18IB, both with an R^2^_p_ of 0.63. C-18IB also had the least favorable RMSE_p_ at 3.55. Squared Spearman’s rank correlation (R^2^_sp_) values (0.57 - 0.89) were similar to R^2^_p_ (0.62 - 0.89) in most trials. The optimal number of components in the most iterations of subsampling, ranged from 2-10, again showing no identifiable pattern according to year, location, population, or sample preparation method (Table 3). Results showing the same general trends with 12 different preprocessing technique combinations for PLSR as well as RF and SVM can be found in Supplemental Table S2.

**Table 3.**
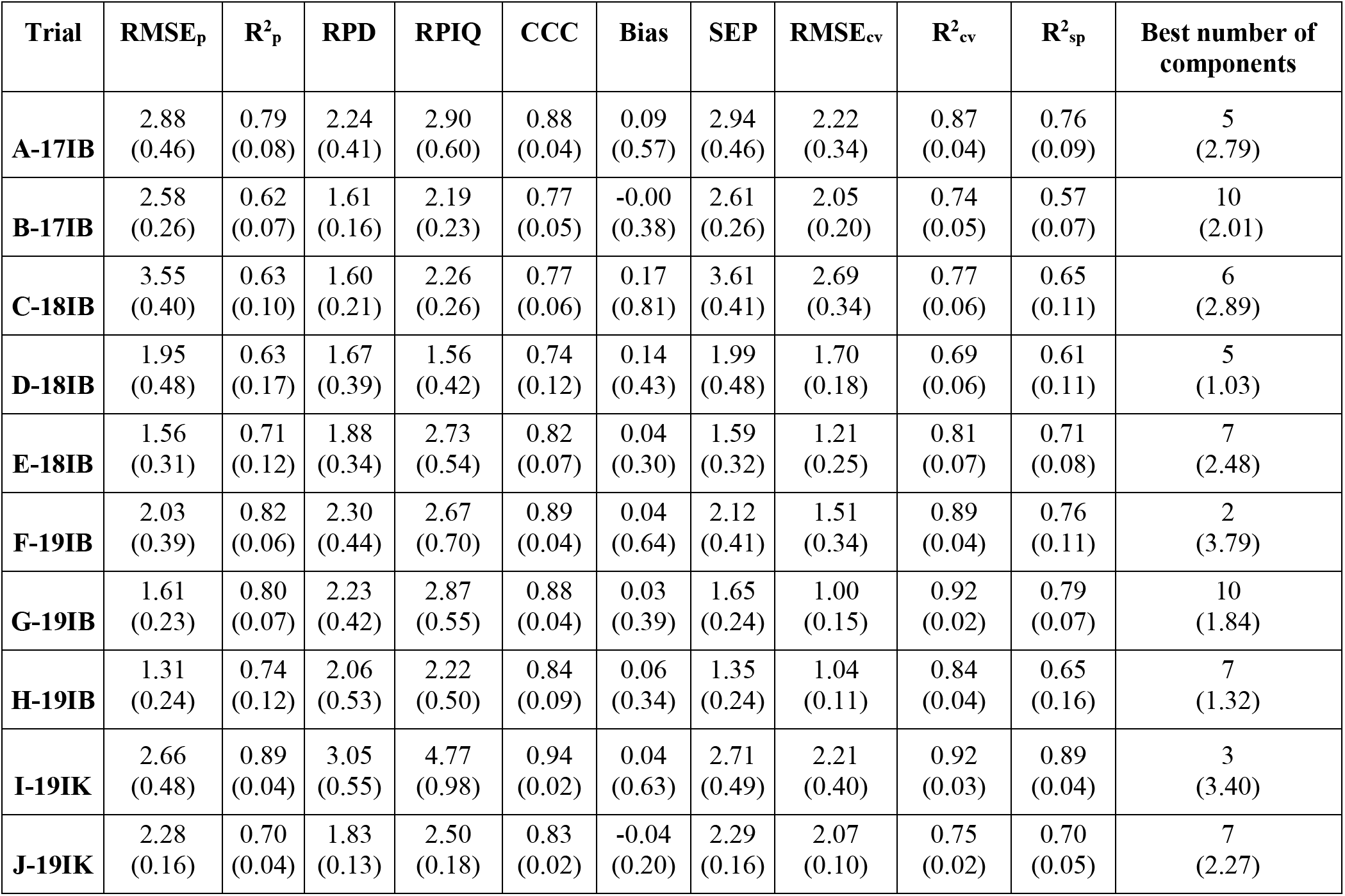
Summary statistics for within-trial cassava root dry matter content partial least squares regression predictions with 50 iterations of subsampling. The mean of each statistic over all iterations is shown for raw data post-outlier removal but without preprocessing except for the number of components, in which the mode is displayed. The standard deviation for each statistic is shown in parentheses. Model performance statistics include: Lin’s concordance correlation coefficient (CCC), coefficient of multiple determination of cross-validation (R^2^_cv_), squared Pearson’s correlation of predicted and observed values in a test set (R^2^_p_), squared Spearman’s correlation of predicted and observed values in a test set (R^2^_sp_), root mean squared error of crossvalidation (RMSE_cv_), root mean squared error of prediction (RMSE_p_), residual predictive deviation (RPD), ratio of performance to interquartile distance (RPIQ), and standard error of prediction (SEP).

| Trial | $RMSE_p$ | $R^2_p$ | RPD | RPIQ | CCC | Bias | SEP | $RMSE_{cv}$ | $R^2_{cv}$ | $R^2_{sp}$ | Best number of components |
| --- | --- | --- | --- | --- | --- | --- | --- | --- | --- | --- | --- |
| **A-17IB** | 2.88<br>(0.46) | 0.79<br>(0.08) | 2.24<br>(0.41) | 2.90<br>(0.60) | 0.88<br>(0.04) | 0.09<br>(0.57) | 2.94<br>(0.46) | 2.22<br>(0.34) | 0.87<br>(0.04) | 0.76<br>(0.09) | 5<br>(2.79) |
| **B-17IB** | 2.58<br>(0.26) | 0.62<br>(0.07) | 1.61<br>(0.16) | 2.19<br>(0.23) | 0.77<br>(0.05) | -0.00<br>(0.38) | 2.61<br>(0.26) | 2.05<br>(0.20) | 0.74<br>(0.05) | 0.57<br>(0.07) | 10<br>(2.01) |
| **C-18IB** | 3.55<br>(0.40) | 0.63<br>(0.10) | 1.60<br>(0.21) | 2.26<br>(0.26) | 0.77<br>(0.06) | 0.17<br>(0.81) | 3.61<br>(0.41) | 2.69<br>(0.34) | 0.77<br>(0.06) | 0.65<br>(0.11) | 6<br>(2.89) |
| **D-18IB** | 1.95<br>(0.48) | 0.63<br>(0.17) | 1.67<br>(0.39) | 1.56<br>(0.42) | 0.74<br>(0.12) | 0.14<br>(0.43) | 1.99<br>(0.48) | 1.70<br>(0.18) | 0.69<br>(0.06) | 0.61<br>(0.11) | 5<br>(1.03) |
| **E-18IB** | 1.56<br>(0.31) | 0.71<br>(0.12) | 1.88<br>(0.34) | 2.73<br>(0.54) | 0.82<br>(0.07) | 0.04<br>(0.30) | 1.59<br>(0.32) | 1.21<br>(0.25) | 0.81<br>(0.07) | 0.71<br>(0.08) | 7<br>(2.48) |
| **F-19IB** | 2.03<br>(0.39) | 0.82<br>(0.06) | 2.30<br>(0.44) | 2.67<br>(0.70) | 0.89<br>(0.04) | 0.04<br>(0.64) | 2.12<br>(0.41) | 1.51<br>(0.34) | 0.89<br>(0.04) | 0.76<br>(0.11) | 2<br>(3.79) |
| **G-19IB** | 1.61<br>(0.23) | 0.80<br>(0.07) | 2.23<br>(0.42) | 2.87<br>(0.55) | 0.88<br>(0.04) | 0.03<br>(0.39) | 1.65<br>(0.24) | 1.00<br>(0.15) | 0.92<br>(0.02) | 0.79<br>(0.07) | 10<br>(1.84) |
| **H-19IB** | 1.31<br>(0.24) | 0.74<br>(0.12) | 2.06<br>(0.53) | 2.22<br>(0.50) | 0.84<br>(0.09) | 0.06<br>(0.34) | 1.35<br>(0.24) | 1.04<br>(0.11) | 0.84<br>(0.04) | 0.65<br>(0.16) | 7<br>(1.32) |
| **I-19IK** | 2.66<br>(0.48) | 0.89<br>(0.04) | 3.05<br>(0.55) | 4.77<br>(0.98) | 0.94<br>(0.02) | 0.04<br>(0.63) | 2.71<br>(0.49) | 2.21<br>(0.40) | 0.92<br>(0.03) | 0.89<br>(0.04) | 3<br>(3.40) |
| **J-19IK** | 2.28<br>(0.16) | 0.70<br>(0.04) | 1.83<br>(0.13) | 2.50<br>(0.18) | 0.83<br>(0.02) | -0.04<br>(0.20) | 2.29<br>(0.16) | 2.07<br>(0.10) | 0.75<br>(0.02) | 0.70<br>(0.05) | 7<br>(2.27) |

**Figure 4.**
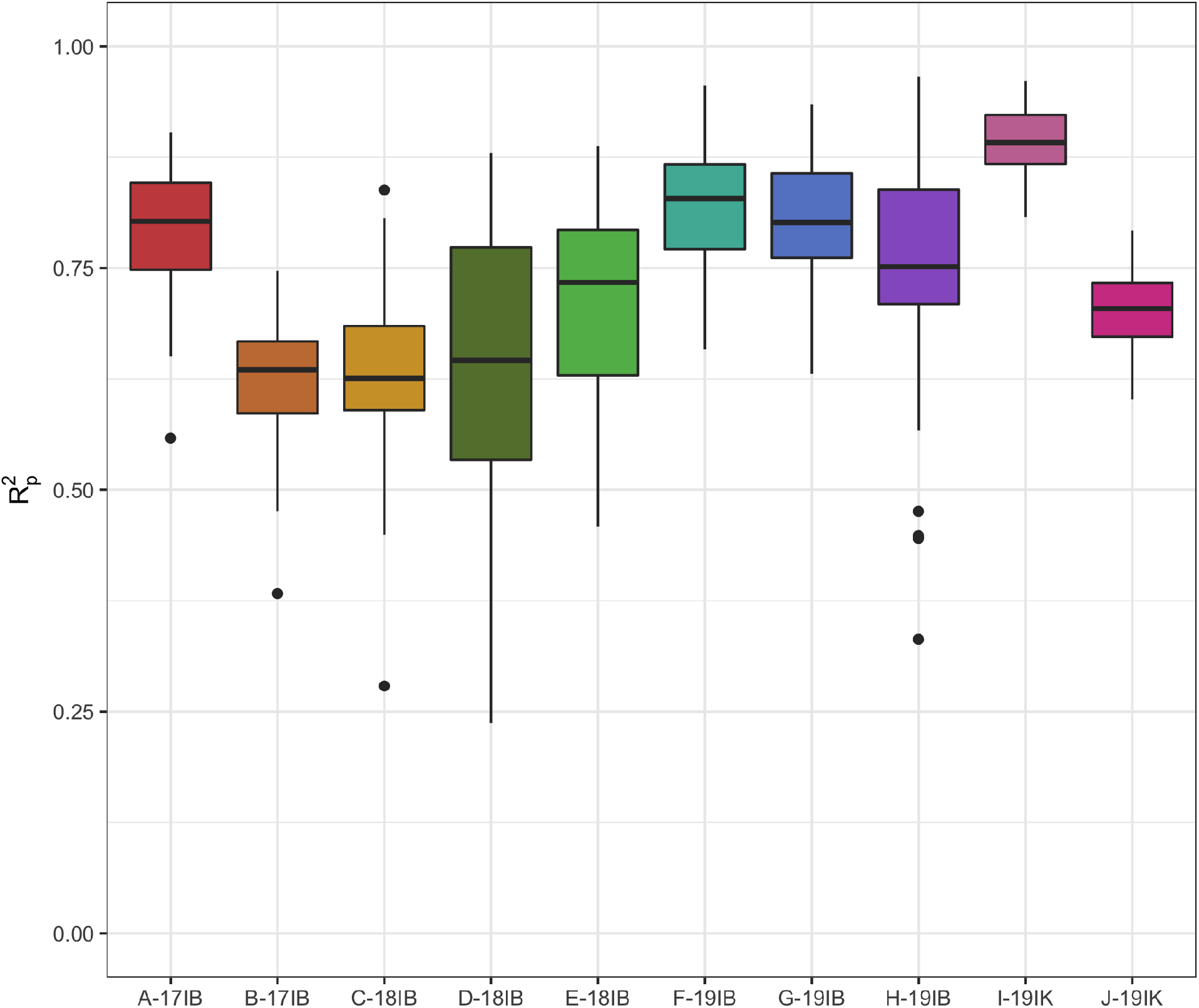
Within-trial predictions of cassava root dry matter content were performed on a plot basis for each trial. R^2^_p_ (squared Pearson’s correlation between predicted and observed root dry matter content) is shown for 50 iterations of the *waves* prediction pipeline with no spectral pretreatment.

Statistics reported in terms of prediction of a holdout test set (e.g., R^2^_p_ and RMSE_p_) show weaker model performance as compared to those reported in terms of CV (R^2^_cv_ and RMSE_cv_) across all trials in this study. In previous studies of NIRS for RDMC prediction in cassava, only leave-one-out CV statistics are discussed, though R^2^_p_ statistics are also reported (Ikeogu et al., 2017; Sánchez et al., 2014). This can lead to misleading conclusions, as we can expect the predictions of future samples to perform more similarly to the R^2^_p_ results rather than those of R^2^_cv_ due to the new samples not being present in the training set. When we directly compare our R^2^_p_ values to those found in previous studies, the performance of the laboratory-grade FOSS 6500 (R^2^_p_ = 0.79 - 0.95) and QualitySpec Trek (R^2^_p_ = 0.84) spectrometers are very comparable to our R^2^_p_ results with the SCiO (within-trial PLSR R^2^_p_ = 0.63 - 0.89), indicating that the SCiO may be a suitable replacement for either of these more expensive spectrometers for RDMC phenotyping, especially when factoring in cost and throughput.

### Algorithms and preprocessing

Many spectral preprocessing methods have been explored (reviewed in Rinnan, van den Berg, & Engelsen, 2009) to reduce noise, but the benefit of individual methods varies depending on the trait of interest, sample set, and model algorithm (Ikeogu et al., 2017; Kosmowski & Worku, 2018; Pizarro et al., 2004). Because the SCiO had not yet been tested for use in the prediction of cassava RDMC, we tested 12 combinations of preprocessing techniques and three model algorithms to identify the combination that provided the best model performance for this spectrometer. Overall, we found no additional benefit to machine learning algorithms as opposed to the more traditional PLSR method (Supplemental Table S2). Though working with group discrimination rather than regression, Kosmowski and Worku (2018) also found that machine learning algorithms were not superior to PLS when investigating SCiO performance. Interestingly, they also found that preprocessing techniques did not impact model performance with PLS or SVM but that data pretreatment was necessary to improve performance of RF models. Studies exploring the impact of these factors on model performance with other spectrometers have not followed this same pattern (Ikeogu et al., 2017; Sampaio et al., 2018), indicating that the ideal combination of spectral and model algorithm may be spectrometer specific. This has repercussions for the development of analysis protocols for future spectrometers, as all combinations of these factors must be tested with each additional hardware specification.

### Sample preparation

Cassava RDMC can vary significantly within roots from proximal to distal end (Chávez et al., 2008), so root tissue is traditionally shredded and mixed to get a representative sample for analysis. Because this method requires the time-consuming transport of bulky roots to a laboratory environment, our study included scans of both shredded tissue and roots prepared for scanning with an alternate sliced method that can be performed directly in the field. Comparing results from plot-mean scans, we observed slightly more favorable model performance statistics from trials in which roots were shredded prior to scanning; the mean within-trial R^2^_p_ ranged from 0.62 to 0.79 for the three sliced root trials and 0.63 to 0.89 for the seven shredded root trials (Table 3). These findings align with those of Ikeogu et al. (2017), as they also found scans of shredded root samples to slightly outperform those of sliced roots in terms of cassava RDMC predictions.

The small sample size in this study limits our ability to confidently draw conclusions based on plot-mean scan model performance alone, so we also sought to determine whether our samples captured the plot-level RDMC within each trial. By varying the number of homogenized subsamples or subsets of sliced roots (hereafter samples) included in the plot mean scan, we were able to identify the point at which within-trial prediction model performance began to level off, indicating that additional samples would not further increase performance and therefore within-plot variation had been adequately captured. Despite differences in the minimum number of homogenized subsamples and sets of sliced roots required to stabilize model performance, five samples were sufficient to capture variation in all trials, regardless of sample preparation method (Figure 5). In seven of the 10 trials in this study, a boost in predictive performance occurred as additional samples were included in the mean scan used for model development up to the leveling point. This increase in performance with additional samples is not surprising, as we expect the addition of technical replicates to stabilize measurements within a given experimental unit by controlling measurement error (Blainey et al., 2014). These results can also be used to inform best practices for future scanning protocol development, allowing breeding programs to evaluate whether the potential small gain in model performance is worth the added cost of root transport for homogenization prior to shredding and to choose the number of samples to include when scanning.

**Figure 5.**
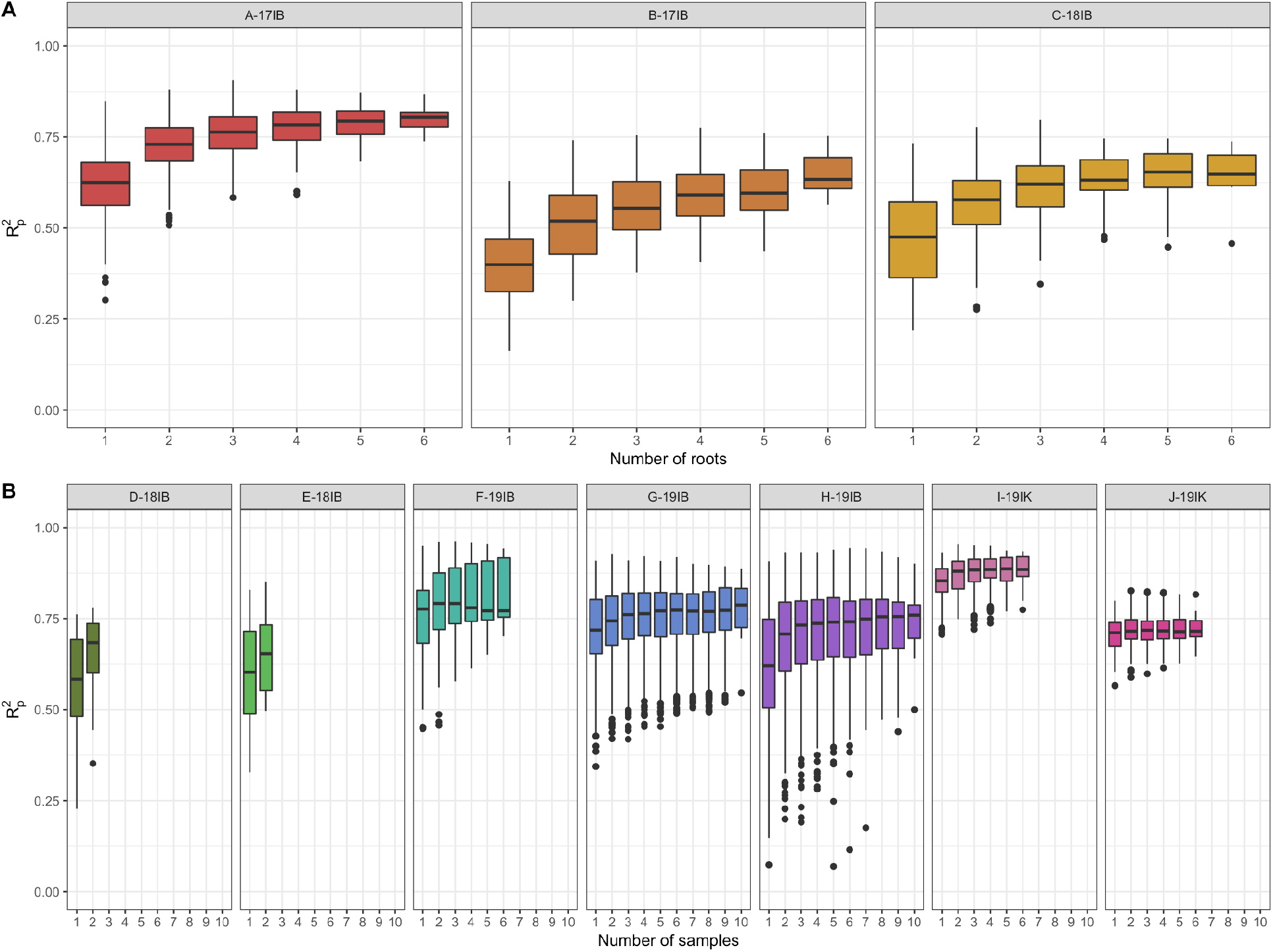
Within-trial predictions of cassava root dry matter content were performed with either subsets of roots (a) or homogenized subsamples (b) for each trial. R^2^_p_ (squared Pearson’s correlation between predicted and observed plot mean dry matter content) is shown for 50 iterations of subsampling of scans within each plot followed by 100 iterations of the *waves* prediction pipeline. The plot-level mean of all scans was taken prior to model development in all cases.

### Biological interpretation of wavelengths

Chemical bonds common in biological tissues (O-H, C-H, N-H) exhibit wavelength-dependent absorption of light in the NIR region, but overlapping signals from complex molecules can be difficult to interpret. In this study, RF variable importance analysis was used to facilitate the biological interpretation of complex spectral signals by ranking the importance of each wavelength in the prediction of cassava RDMC (Figure 6). All trials followed the same general pattern of variable importance, with higher importance measures present on both ends of the spectral range (approximately 740-760 nm and 1050-1070 nm) and around 950-990 nm, an overtone for the oxygen-hydrogen bond and one of the most prominent signals for water in the NIR range. This region has been suggested for use in the prediction of water status in the remote sensing of plants (Peñuelas et al., 1993; Sims & Gamon, 2003). Spectral reflectance at 970 nm has since been included in several vegetation indices for the detection of water status that have been shown to have high correlations with water content in foliage [e.g., water band index (Peñuelas et al., 1993), reflectance water index (Peñuelas et al., 1997), and normalized water indices (Babar et al., 2006; Prasad et al., 2007)].

**Figure 6.**
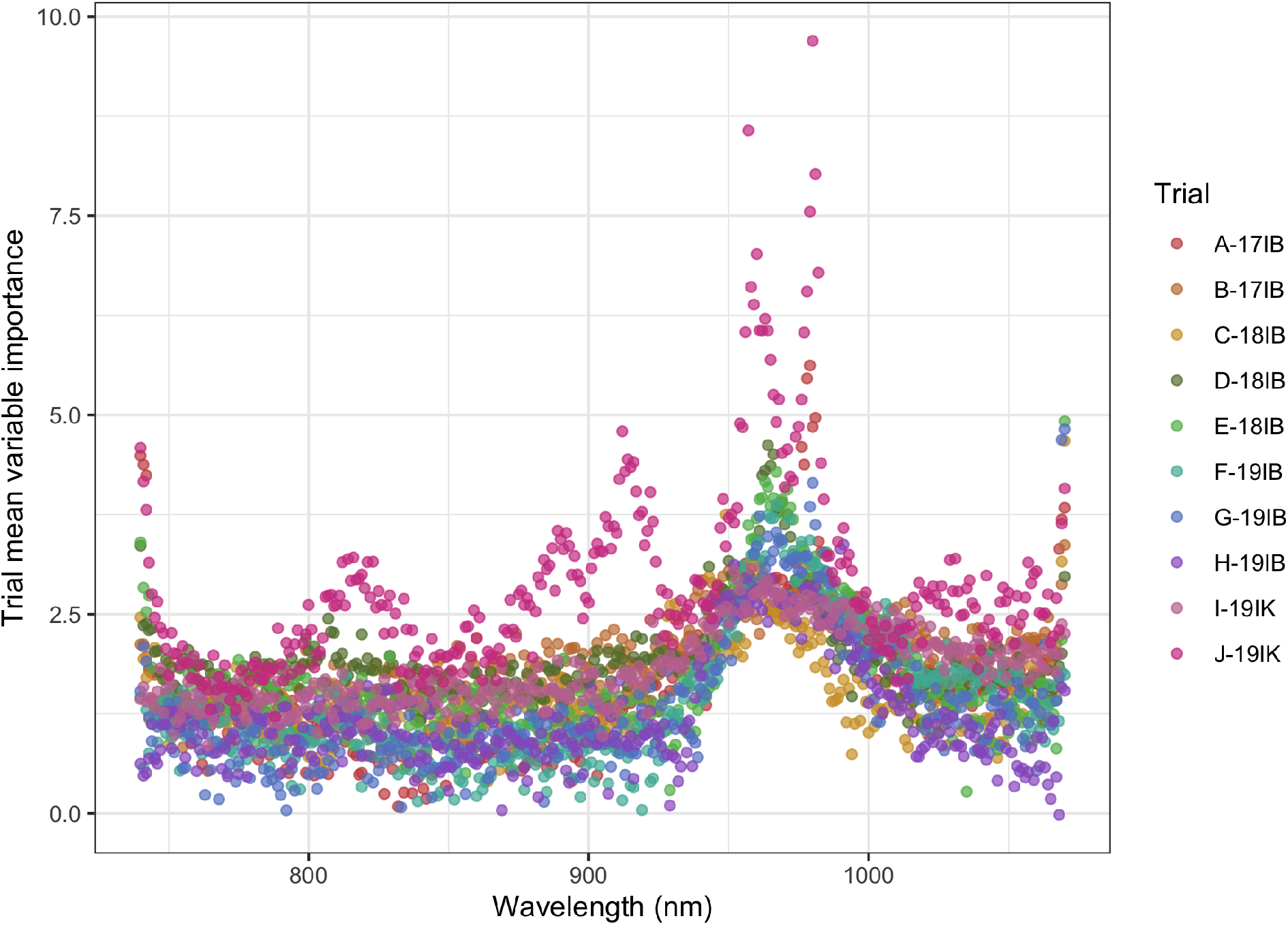
Random forest variable importance analysis was performed for each wavelength on a within-trial basis 10 times. The mean importance value for each wavelength within a trial is represented by a single point.

The presence of this prominent water-associated signal within the range of the SCiO is likely responsible for the successful cassava RDMC predictions we reported in this study. Scans from the SCiO have also been found to be highly predictive in many other plant systems, including seed-based cultivar identification for chickpea, sorghum, and millet (Kosmowski & Worku, 2018), total antioxidant capacity in various gluten-free grains (R^2^_p_ = 0.85 - 0.90) (Wiedemair & Huck, 2018), soluble solids in kiwi, apple maturity, avocado ripeness (Li et al., 2018), and apple (R^2^_p_ = 0.86) and kiwi (R^2^_p_ = 0.84) fruit dry matter content (Kaur et al., 2017). This indicates that the SCiO may be useful in additional traits both within and beyond cassava.

### Application of low-cost NIRS in a cassava breeding program

Optimal NIRS model calibration requires the training set to be representative of the greater population of samples that will be predicted. Because the RDMC of cassava is affected by the environment in which it is grown, we chose to include trials from a range of years, populations, and locations, and to investigate the influence of these factors on model performance using CV schemes designed for plant breeding programs. Overall mean model performance using these schemes ranged from R^2^_p_ = 0.60 with CV0 and CV00 to R^2^_p_ = 0.71 for CV1 (Supplemental Table 4). In general, we found that schemes in which the test set environment was represented in the training set (CV1 and CV2) outperformed schemes in which there was no overlap in environments between the two sets (CV0 and CV00), while the differences in performance within each of these groups was minimal (Figure 7, Supplemental Table 4). This indicates that sharing an environment between the training and test sets has more of an impact on model performance than sharing a set of clones. We therefore recommend that training sets be updated to include a subset of oven-phenotyped plots from each new trial to maximize model performance, as is done in the CV1 and CV2 schemes. However, as the addition of oven-dried phenotyping represents a tradeoff between model performance and labor, each breeding program will need to evaluate whether the additional performance boost is worthwhile. We also observed differences in overall model performance between trials, following the same pattern as our within-trial prediction results. Given the mix of overall trial performance amongst locations, years, populations, and scanning protocols, it is not possible to determine the source of this pattern with the limited sample size of this study. Further investigation into the effects of these factors on RDMC prediction with NIRS would help to clarify these relationships.

**Figure 7.**
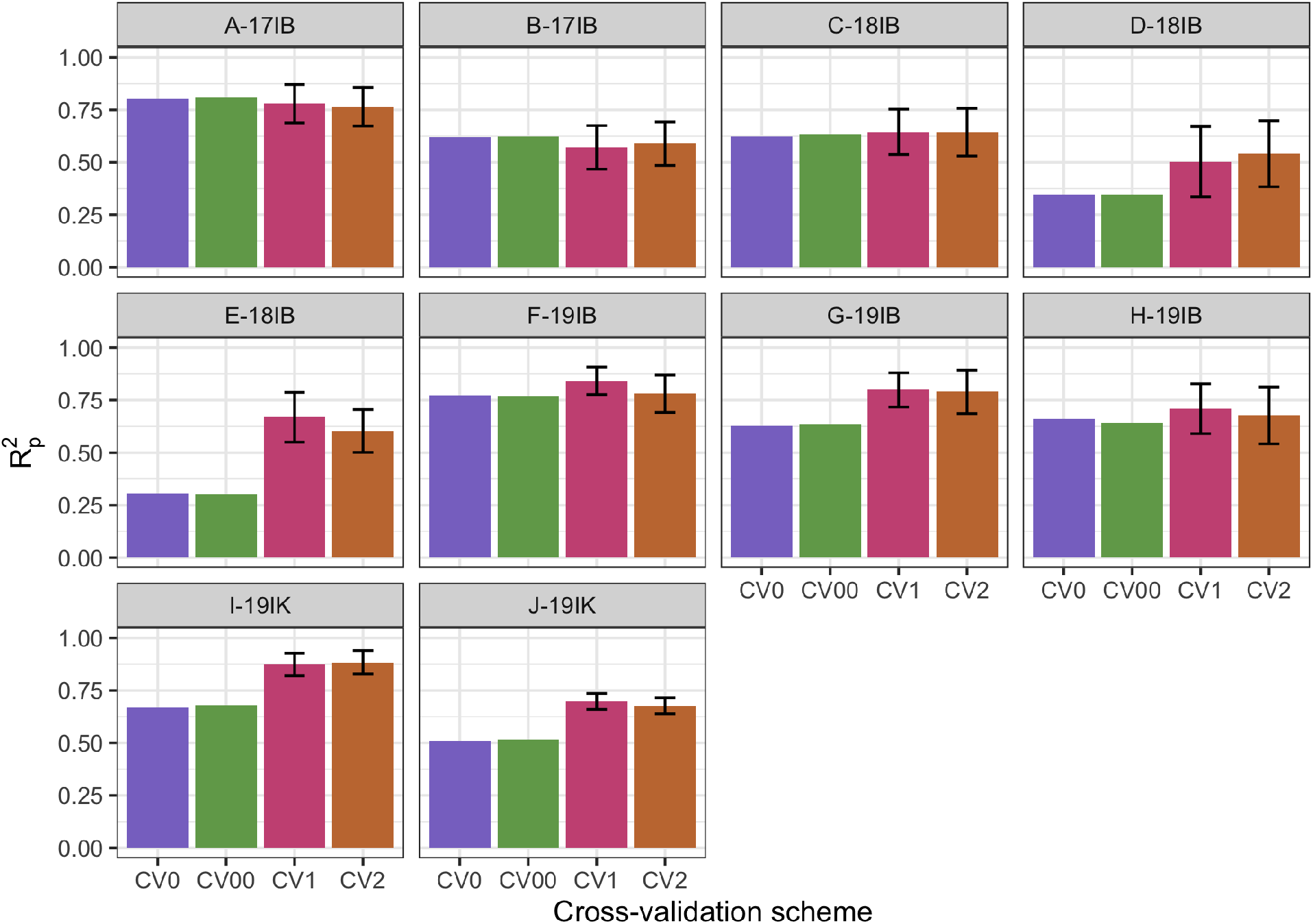
Partial least squares regression prediction of cassava root dry matter content performed on a plot-mean basis for each trial. Cross-validation (CV) scheme results are displayed as the R^2^_p_ (squared Pearson’s correlation between predicted and observed values) for 50 iterations of the *waves* prediction pipeline with no spectral pretreatment. CV0 indicates leave-one-trial-out crossvalidation, CV00 indicates that there was no overlap between clones and environments in the training and test sets, CV1 indicates overlap in environment but not clones between the training and test sets, and CV2 indicates overlap of both clones and environments in the training and test sets, though in all cases clones with multiple replicates within a trial were sorted together. Error bars show standard deviation for schemes with subsampling (CV1 and CV2). As no subsampling occurred in either the CV0 or CV00 schemes, standard deviation was not calculated and therefore no error bars are displayed.

## CONCLUSIONS

The present study shows that miniature NIR spectrometers such as the SCiO have the potential to enable cheap and high-throughput quality assessments particularly for breeding programs that cannot afford expensive, lab-based NIR equipment. The results of this study support the application of the SCiO for the phenotyping of cassava RDMC directly in the field, with model performance matching that of laboratory-grade spectrometers in many trials. The low per-unit cost of this new generation of consumer-grade spectrometers will contribute not only to the efficiency with which cassava breeders are able to phenotype quality traits but also for other starchy root and tuber crops.

Our results indicate that scans taken on sliced roots may match the performance of scans taken on homogenized root subsamples, but additional studies directly comparing these two methods within the same trials would help to clarify this relationship. The ability to accurately predict RDMC with sliced roots represents a large savings in terms of time and labor on the part of breeding programs, allowing maximum benefit from the use of handheld spectrometers. Further, we observed diminishing returns with the inclusion of additional samples per plot, indicating that scanning a subset of five or more roots or alternatively two or more homogenized subsamples should be sufficient for capturing within-plot scan variation with the SCiO. Overall, we found prediction with the SCiO, a consumer-grade NIR spectrometer, to be a highly accurate, field-based, and low-cost method for cassava RDMC phenotyping. The scanning and analysis protocols explored in this study are readily applicable in the phenotyping of other quality traits in cassava and beyond.

## Abbreviations

CV: Cross-validation
R^2^_cv_: Coefficient of multiple determination of cross-validation
IITA: International Institute of Tropical Agriculture
NIRS: Near-infrared spectroscopy
PLSR: Partial least squares regression
RF: Random forest
RMSE_cv_, RDMC: Root dry matter content; Root mean squared error of cross-validation
RMSE_p_: Root mean squared error of prediction
R^2^_p_: Squared Pearson’s correlation between predicted and observed values in a test set
R^2^_sp_: Squared Spearman’s rank correlation between predicted and observed values in a test set
SVM: Support vector machine
Vis: Visible light

## ACKNOWLEDGEMENTS

We are grateful to the Next Generation Cassava team and in particular the IITA cassava breeding team for their interest in and support of this work. This project would not have been possible without their efforts. We also thank the PhenoApps team, Kelly Robbins, Jean-Luc Jannink, and the Gore research group for valuable feedback and productive discussions. This work has been supported by USDA NIFA AFRI EWD Predoctoral Fellowship 2019-67011-29606 (J.H.) and NSF BREAD IOS-1543958 (M.A.G.). Work at IITA was supported by the United Kingdom Foreign, Commonwealth & Development Office (FCDO) and the Bill & Melinda Gates Foundation (Grant INV-007637). This study was also made possible by the support of the American People provided to the Feed the Future Innovation Lab for Crop Improvement through the United States Agency for International Development (USAID). The contents are the sole responsibility of the authors and do not necessarily reflect the views of USAID or the United States Government. Program activities are funded by the United States Agency for International Development (USAID) under Cooperative Agreement No. 7200AA-19LE-00005.

## SUPPLEMENTAL MATERIAL

**Supplemental Figure S1.** Illustrated sample preparation, scanning, and root dry matter content determination protocol

**Supplemental Figure S2.** Within-trial algorithm and preprocessing technique comparison

**Supplemental Table S1.** Spectral outliers

**Supplemental Table S2**. Within-trial prediction model performance summary statistics

**Supplemental Table S3.** Common clone counts

**Supplemental Table S4**. Cross-validation scheme model performance summary statistics

## CONFLICT OF INTEREST

The authors declare no conflict of interest.

## DATA AVAILABILITY

Raw RDMC and spectral data are available for download on Cyverse at https://datacommons.cyverse.org/browse/iplant/home/shared/GoreLab/dataFromPubs/Hershberger_CassavaNIRS_2021/. Analysis R code is available on Github at www.github.com/GoreLab/CassavaNIRS.

## AUTHOR CONTRIBUTIONS

JH - conceptualization, methodology development, writing of original draft, data curation, formal analysis, revision and editing

EGNM - data collection, revision and editing

PP - methodology development, data collection, data curation

ASI - methodology development, data collection

KO - methodology development, data collection

KN - methodology development, data collection

IYR - experimental design, supervision, funding acquisition, revision and editing

MAG - conceptualization, supervision, funding acquisition, revision and editing

## Notes

https://www.github.com/GoreLab/CassavaNIRS

https://datacommons.cyverse.org/browse/iplant/home/shared/GoreLab/dataFromPubs/Hershberger_CassavaNIRS_2021/

### Competing Interest Statement

The authors have declared no competing interest.

