## Supplemental Tables and Figures for "Low-cost, handheld near-infrared spectroscopy for root dry matter content prediction in cassava"

**SUPPLEMENTAL MATERIAL**

*Supplemental Figures*
S1. Illustrated sample preparation, scanning, and root dry matter content determination protocol
S2. Within-trial algorithm and preprocessing technique comparison
*Supplemental Tables*
S1. Spectral outliers
S2. Within-trial prediction model performance summary statistics
S3. Common clone counts
S4. Cross-validation scheme model performance summary statistics

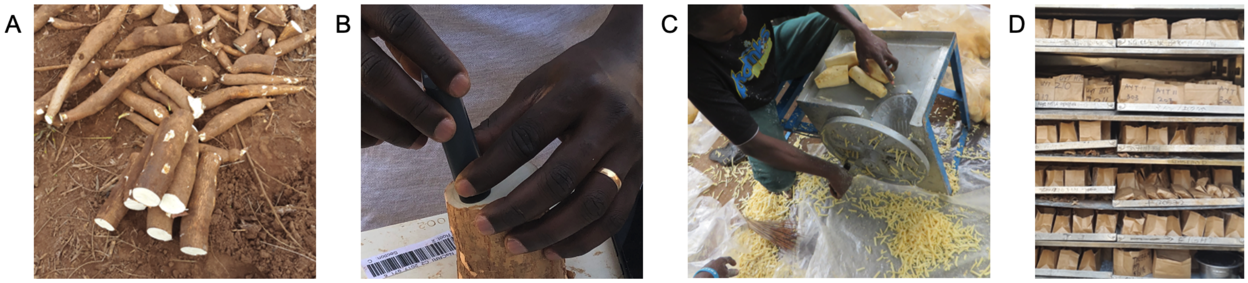


**Figure S1. Illustrated sample preparation, scanning, and root dry matter content determination protocol.** Six commercial-sized cassava roots were harvested per plot (A). Roots from trials A-17IB, B-17IB, and C-18IB were sliced crosswise in the center and the cut surface was scanned as in (B). Scanned roots were then shredded (C) for dry matter content determination using the oven method (D). Roots from all other trials were homogenized by shredding (C) prior to scanning and dry matter determination.

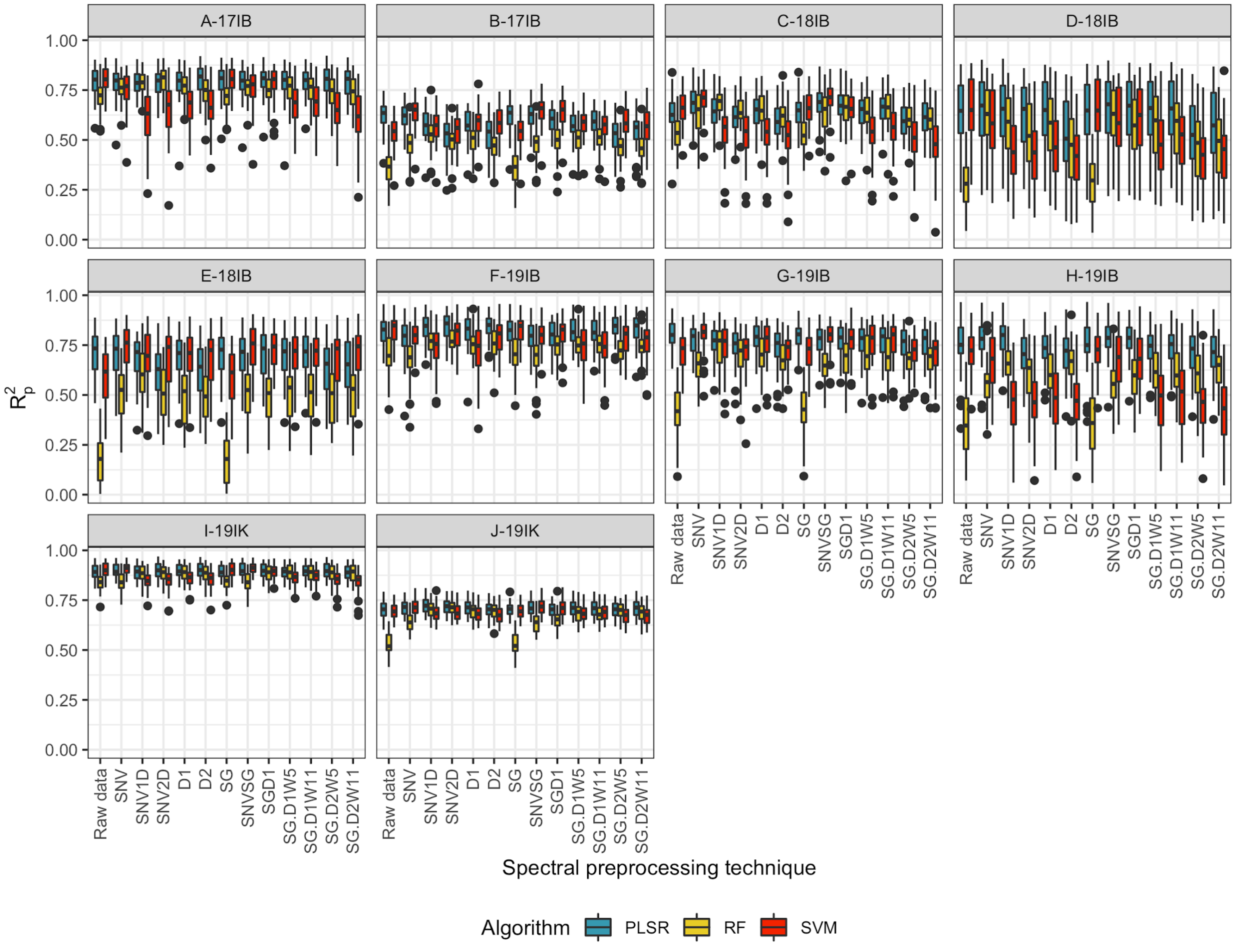


**Figure S2. Within-trial algorithm and preprocessing technique comparison.** Within-trial prediction of cassava root dry matter content was performed on a plot-basis for each trial. Model performance (squared Pearson's correlation between predicted and observed values; $R^2{}_p$) is displayed for 50 iterations of the waves prediction pipeline in R. Results are grouped by trial with an algorithm represented by color and spectral preprocessing technique on the x-axis. In addition to models without spectral preprocessing (raw data), preprocessing techniques include standard normal variate (SNV), standard normal variate and first derivative (SNV1D), standard normal variate and second derivative (SNV2D), first derivative (D1), second derivative (D2), Savitzky-Golay with window size = 11 (SG), standard normal variate and Savitzky-Golay (SNVSG), gap segment derivative with window size = 11 (SGD1), Savitzky-Golay with window size = 5 and first derivative (SG.D1W5), Savitzky-Golay with window size = 11 and first derivative (SG.D1W11), Savitzky-Golay with window size = 5 and second derivative (SG.D2W5), and Savitzky-Golay with window size = 11 and second derivative (SG.D2W11).

**Table S1. Spectral outliers.** Metadata for outlier scans as identified through Mahalanobis distance exceeding the outlier threshold as determined by a $\chi^2$-distribution with 331 degrees of freedom.

| Trial | Plot Number | Root Number | Subsample | Scan Timestamp | Device ID | Mahalanobis Distance |
|---|---|---|---|---|---|---|
| C-18IB | 138 | 3 | 3 | 6/18/19 | 9026A17D517 9A429 | 22283.604 |
| G-19IB | 210 | NA | 10 | 4/27/20 | D032D9BF4E 067612 | 22155.2747 |
| A-17IB | 1007 | 3 | 3 | 4/23/18 | 9026A17D517 9A429 | 4303.80184 |
| A-17IB | 1005 | 6 | 6 | 4/23/18 | 9026A17D517 9A429 | 867.104283 |
| C-18IB | 125 | 4 | 4 | 6/18/19 | 9026A17D517 9A429 | 809.519396 |
| A-17IB | 1003 | 4 | 4 | 4/23/18 | 9026A17D517 9A429 | 779.010815 |
| A-17IB | 2030 | 5 | 5 | 5/8/18 | D032D9BF4E 067612 | 518.599427 |
| E-18IB | 332 | NA | 2 | 9/23/19 | D032D9BF4E 067612 | 469.645697 |

**Table S2. Within-trial prediction model performance summary statistics.** Within-trial model performance summary statistics for partial least squares regression (PLSR), random forest (RF), and support vector machine (SVM) algorithms with raw data and 12 preprocessing techniques applied before model fitting. Model performance statistics include: Lin's concordance correlation coefficient (CCC), coefficient of multiple determination of cross-validation ($R^2_{cv}$), squared Pearson's correlation of predicted and observed values in a test set ($R^2_p$), squared Spearman's correlation of predicted and observed values in a test set ($R^2_{sp}$), root mean squared error of cross-validation ($RMSE_{cv}$), root mean squared error of prediction ($RMSE_p$), residual predictive deviation (RPD), ratio of performance to interquartile distance (RPIQ), and standard error of prediction (SEP). Spectral pretreatments include: standard normal variate (SNV), standard normal variate and first derivative (SNV1D), standard normal variate and second derivative (SNV2D), first derivative (D1), second derivative (D2), Savitzky-Golay with window size = 11 (SG), standard normal variate and Savitzky-Golay (SNVSG), gap segment derivative with window size = 11 (SGD1), Savitzky-Golay with window size = 5 and first derivative (SG.D1W5), Savitzky-Golay with window size = 11 and first derivative (SG.D1W11), Savitzky-Golay with window size = 5 and second derivative (SG.D2W5), and Savitzky-Golay with window size = 11 and second derivative (SG.D2W11).

Due to the large size of this table, it is available in the file "Supplemental_Table_S2_within_trial_predictions.csv" and can be downloaded from Cyverse: https://datacommons.cyverse.org/browse/iplant/home/shared/GoreLab/dataFromPubs/Hershberger_CassavaNIRS_2021/Supplemental_files/

**Table S3. Common clone counts.** Counts of unique clones that were phenotyped in each trial are shown in bold on the diagonal. The number of phenotyped clones in common between two trials is shown on the off-diagonals.

| Trial | A-17IB | B-17IB | C-18IB | D-18IB | E-18IB | F-19IB | G-19IB | H-19IB | I-19IK | J-19IK |
|---|---|---|---|---|---|---|---|---|---|---|
| **A-17IB** | **48** | 4 | 40 | 4 | 4 | 4 | 4 | 4 | 39 | 3 |
| **B-17IB** | 4 | **65** | 5 | 5 | 5 | 4 | 5 | 5 | 5 | 4 |
| **C-18IB** | 40 | 5 | **51** | 7 | 10 | 10 | 5 | 5 | 50 | 5 |
| **D-18IB** | 4 | 5 | 7 | **35** | 7 | 6 | 33 | 5 | 7 | 5 |
| **E-18IB** | 4 | 5 | 10 | 7 | **36** | 8 | 5 | 29 | 10 | 5 |
| **F-19IB** | 4 | 4 | 10 | 6 | 8 | **30** | 4 | 4 | 10 | 6 |
| **G-19IB** | 4 | 5 | 5 | 33 | 5 | 4 | **35** | 5 | 5 | 4 |
| **H-19IB** | 4 | 5 | 5 | 5 | 29 | 4 | 5 | **36** | 5 | 4 |
| **I-19IK** | 39 | 5 | 50 | 7 | 10 | 10 | 5 | 5 | **50** | 5 |
| **J-19IK** | 3 | 4 | 5 | 5 | 5 | 6 | 4 | 4 | 5 | **180** |

**Table S4. Cross-validation scheme model performance summary statistics.** Model performance summary statistics for partial least squares regression (PLSR) using raw spectral data under four plant breeding cross-validation (CV) schemes. CV0 indicates leave-one-trial out cross-validation, CV00 indicates that there was no overlap between clones and environments in the training and test sets, CV1 indicates overlap in environment but not clones between the training and test sets, and CV2 indicates overlap of both clones and environments in the training and test sets, though in all cases clones with multiple replicates within a trial were sorted together. Model performance statistics include: Lin's concordance correlation coefficient (CCC), coefficient of multiple determination of cross-validation ($R^2_{cv}$), squared Pearson's correlation of predicted and observed values in a test set ($R^2_p$), squared Spearman's correlation of predicted and observed values in a test set ($R^2_{sp}$), root mean squared error of cross-validation ($RMSE_{cv}$), root mean squared error of prediction ($RMSE_p$), residual predictive deviation (RPD), ratio of performance to interquartile distance (RPIQ), and standard error of prediction (SEP).

Due to the large size of this table, it is available in the file "Supplemental_Table_S4_cv_results.csv" and can be downloaded from Cyverse:

https://datacommons.cyverse.org/browse/iplant/home/shared/GoreLab/dataFromPubs/Hershberger_CassavaNIRS_2021/Supplemental_files/
